# Genetic similarity and facial cues for kin recognition in humans

**DOI:** 10.1101/2020.04.06.028134

**Authors:** Nicole S. Torosin, Joshua Ward, Adrian V. Bell, Leslie A. Knapp

## Abstract

Kin recognition is essential to the evolution of human cooperation, social organization, and altruistic behavior. However, the genetic underpinnings of kin recognition have been largely understudied. Facial resemblance is an important relatedness cue for humans and more closely related individuals are generally thought to share greater facial similarity. To evaluate the relationship between perceived self-resemblance and genetic similarity among biologically related and unrelated females, we administered facial self-recognition surveys to twenty-three sets of related females and genotyped three different genetic systems, human leukocyte antigens (HLA), neutral nuclear microsatellites and mitochondrial haplogroups, for each individual. Using these data, we examined the relationship between visual kin recognition and genetic similarity. We found that pairs of individuals identified as visually more similar had greater HLA allelic sharing when compared to less facially similar participants. We did not find the same relationship for microsatellite and mitochondrial similarity, suggesting that HLA allelic similarity increases the probability of perceived self-resemblance in humans while other genetic markers do not. Our results demonstrate that some genetic markers, such as HLA-DRB, may have significant influence on phenotype and that large scale surveys of HLA and facial feature morphology will yield valuable insight into the evolutionary biology of genotype-phenotype relationships and kin recognition.

## 1 Introduction

Kin recognition, and the ability to differentiate between individuals with different degrees of genetic relatedness, is fundamental to understanding a range of phenomena across taxa. Aside from its relationship to the evolution of cooperation (e.g. [23, 22, 25, 43, 29, 4, 19]), the ability to distinguish kin from non-kin may impact how social kinship is organized, how it evolves, and how mates are selected [32, 30]. However, the genetic underpinnings of kin recognition have been largely understudied.

Experimental studies have found that humans use facial resemblance as a relatedness cue and concomitantly adjust their behavior towards others [14]. There is evidence that self-resemblance leads to increased cooperative behavior [16], more trust [12, 13, 36], a greater likelihood of altruistic behavior [15] and greater paternal investment [2]. Even computer manipulated images are perceived as more trustworthy when the image is similar to the participant [12]. In the present study, we explore how genetic factors, specifically genetic similarity, play a role in providing visual cues of kin recognition and perceptions of facial resemblance in humans.

Research on perceived relatedness in humans has yielded conflicting results. Christenfeld and Hill [11] reported that naïve observers were better able to correctly pair one-year-old children with their fathers than with their mothers. McLain et al. [33] found that judges matched photographs of biological mothers to their newborns significantly more frequently than they matched biological fathers to their newborns. Other studies have not always replicated these results. Some research, instead, found that children are perceived as similar to both their fathers and mothers [7, 8, 34].

It has been suggested that patterns of parent-child phenotypic similarity vary according to the age of the child. For example, younger children are thought to resemble their mothers and older children may have a greater resemblance to their fathers [2]. Alvergne et. al. [3] demonstrated that unrelated judges can recognize parent–child pairs through facial photographs from families belonging to both their own cultural groups and from different origins. Using photographs, unrelated observers are also able to identify pairs of related children [31]. Additionally, same sex pairs of adult siblings can also be identified by unrelated observers but recognition of opposite sex adult siblings appears to be much more difficult[17].

To help clarify the relationship between relatedness and self-resemblance, we examined the relationship between visual kin recognition and genetic similarity using three different genetic systems: human leukocyte antigens (HLA), which are known as the major histocompatibility complex (MHC) in non-human organisms, neutral microsatellites found in the nuclear genome, and mitochondrial haplogroups. The HLA plays a critical role in immune response and regulation. HLA genes are highly polymorphic and although frequently shared by related individuals [6], they typically differ between unrelated individuals. While it is suggested that olfactory kin recognition cues are the consequence of an individual’s HLA genotype [42], the relationship between HLA genotype and visual kin recognition has been unexplored. We hypothesized that individuals identified as the most visually similar to study participants would share more HLA alleles compared to visually dissimilar individuals.

## 2 Methods

### Facial Images

Twenty-three sets of related females living in Oklahoma were recruited for a total of 69 participants. Each related set of three females will henceforth be called a “triad.” The University of Cambridge Human Biology Research Ethics Committee approved our research (Application No. 2008.06) and each participant provided informed consent. All individuals self-identified as Caucasian and all had similar skin tones. To minimize any differences in face coloration, and to standardize background color of all photographs, we took black and white photographs of each participant. Participants removed makeup, facial jewelry and pushed their hair off their foreheads. Photographs were taken from one meter away under identical lighting conditions. Participants assumed neutral expressions with closed mouth and open eyes. We used Adobe Photoshop CS3 for Windows to align the photographs, making each the same dimensions and aligning the eyes on the same x-axis. For all surveys, we cropped the participant’s hair, ears, and neck from each photo and presented photographs of a facial image only.

### Facial Similarity Tests

To assess a participant’s perception of their own facial similarity to kin and non-kin, a Facial Self-Recognition (FSR) test was administered. For the FSR, participants were shown an A2 sized piece of paper with photos of the other 68 faces. Each participant was instructed to choose and then rank, in order of similarity, the three faces they perceived as facially similar to themselves.

### Laboratory methods and genotyping

To evaluate the relationship between perceived facial similarity and genetic similarity, we determined HLA-DRB genotypes, identified alleles for eight highly polymorphic, unlinked, microsatellite loci, and sequenced each participant’s complete mitochondrial D-loop (also known as the hypervariable regions, HVI and HVII). Buccal swabs and hair follicles were collected from all participants and samples were stored at room temperature for approximately one month prior to extraction. A Chelex^®^ extraction protocol was used to obtain DNA from buccal swabs [41]. For two samples, extractions from buccal swabs failed so a standard organic extraction procedure was used with plucked hair samples [39].

We identified HLA-DRB genotypes using polymerase chain reaction (PCR) to amplify the highly variable second exon of all HLA-DRB loci. Denaturing gradient gel electrophoresis (DGGE), followed by direct sequencing, was used to identify precise DRB alleles [27]. All HLA sequences were compared to the IMGT/HLA Database (www.Imgt.org) to determine specific HLA-DRB alleles. We grouped HLA-DRB alleles into eight serotypes, a designation of functional similarity determined by allelic association with the same antibodies [18]. We did not group alleles into HLA supertypes because we found only four different supertypes in our participant population.

Microsatellite (MSAT) loci with high rates of heterozygosity were amplified using PCR in multiplex reactions. Multiplex polymerase chain reactions (PCRs) were prepared using Qiagen Multiplex PCR reagents (Manchester, UK), 60ng BSA (Sigma, Haverhill, U.K.) and 450 nM each forward and reverse primers. We constructed three microsatellite multiplexes: 1) D1s533, D12s375 and D18s537, 2) D2s1326, D5s1457 and D9s9222 and 3) D13s317 and D3s1766. PCR was carried out using a Techne Genius Thermal Cycler with the following cycling conditions: 95*°C* for15 minutes, then 40 cycles of 94*°C* for 30 seconds, 58*°C* for 90 seconds) and 72*°C* for 60 seconds, followed by a final extension of 60*°C* for 30 minutes. We analyzed amplicons using fluorescence and alleles for each locus were identified using PeakScanner2 software (Thermo Fisher, Loughborough, U.K.). Mitochondrial DNA (mtDNA) HVI and HVII regions were PCR amplified and amplicons were directly sequenced to determine each participant’s mitochondrial haplotype [20]. Supplemental **Table S4** reports the HLA-DRB, serotype, MSAT, and mtDNA genotype information for each individual.

### Allele Sharing

To measure allele sharing between participants we used the Degree of Individuality (*D*) index [44].

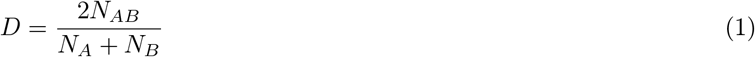

Where *N_AB_* = Total number of alleles (or HLA-DRB serotypes) shared between individuals *A* and *B, N_A_* = total number of alleles (or serotypes) identified in individual *A, N_B_* = Total number of alleles (or serotypes) identified in individual *B*. A *D*-score of zero indicates there were no shared alleles (or serotypes) between individuals and therefore a low chance of relatedness. *A D*-score of one indicates that two individuals share all alleles (or serotypes) and therefore have a high chance of relatedness. *A D*-score of less than 0.25 indicates non-related individuals, while a *D*-score of equal or greater to 0.50 indicates a first-degree relationship (e.g., mother and daughter). Our predictions were that mothers and daughters would possess a *D*-score of 0.50 and that sisters would also have *D*-scores of 0.50.

### Statistical analyses

To assess whether HLA-DRB and/or MSAT allele sharing was associated with perception of selfsimilarity, we fit an exploded logit model. We selected this model because it can predict how different variables will affect ranked data [1]. In this study, participants ranked self-similarity. Variables potentially affecting the ranked data included genetic similarity, relatedness, and age difference. In this statistical framework, each participant *i* has a certain utility *u_ij_* for another individual *j*. Assuming that respondent *i* will give individual *j* a better rank than individual *k* when *u_ij_* > *u_ik_*, we use the utility function to model the predictor variables: 1) the D-score, 2) coefficient of relationship, and 3) the absolute age difference between each individual and the participant. To evaluate how much the genetic D-score altered the probability that the person would be chosen based on phenotypic similarity to the participant, a linear combination of the three variables was used,

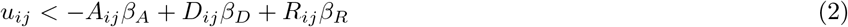

Where *A_ij_* is the absolute value of the age difference, *D_ij_* is the genetic *D*-score, and *R_ij_* the coefficient of relationship between the participant *i* and individual *j*. Values of *R_ij_* are based on the average genetic similarity a pair is predicted to have. For example, mother-daughter: 50%, sisters: 50%, grandmother-granddaughter: 25%. For everyone outside a participant’s triad we assumed 0% relatedness.

The likelihood of the model and the maximum likelihood estimates of *β_A_, β_D_*, and *β_R_* were found through Nelder-Mead optimization. Standard errors were found by estimating the Hessian matrix and solving for the variance-covariance matrix. Three variations of the model were fit and AIC goodness-of-fit scores were computed to compare model performance (**Table S1-S3**). The R code [37] is found in the Supplemental Material.

## 3 Results

Related female triads included seventeen mother-daughter-daughter sets, four sister-sister-sister sets, and two grandmotherdaughter-granddaughter sets. We identified thirteen HLA-DRB alleles from six different HLA-DRB loci (HLA-DRB1, -DRB3, -DRB4, -DRB5 and -DRB6), eight HLA-DRB serotypes, and an average of eight microsatellite alleles for each locus in our 23 triads. Nine mother-daughter pairs were HLA-DRB identical and eleven sister-sister pairs were HLA-DRB identical. We found a relationship between HLA-DRB *D*-scores and MSAT *D*-scores between individuals (R-squared= 0.047, p= 1.45E-51), suggesting that HLA-DRB allelic diversity is correlated with overall genomic diversity as measured by microsatellites (**Figure S1**).

Our Facial Self-Recognition (FSR) tests required each participant to identify faces that they thought were the first, second and third most similar to their own. (For FSR data table see **Table S4**). In all cases, a participant’s first and second choices were the other members of their own triad. The third choice was, by necessity, an unrelated individual.

To test whether HLA-DRB *D*-score between participants in the FSR survey altered the probability of an individual being chosen, we fit the exploded logit model specified above (**Figure 1**). The model incorporated *D*-scores, the coefficient of relationship, and the age difference between each participant and the other 68 possible choices. Figure 1 shows that the coefficient of relationship is an important predictor of choice. Each 0.25 above the average R score increased the probability of an individual being chosen by 0.1. For each 0.1 *D*-score point above average between two individuals, the probability of an individual being chosen increased by a few percentage points. The same trend was observed for HLA-DRB serotypes (**Figure S2**).

**Figure 1:**
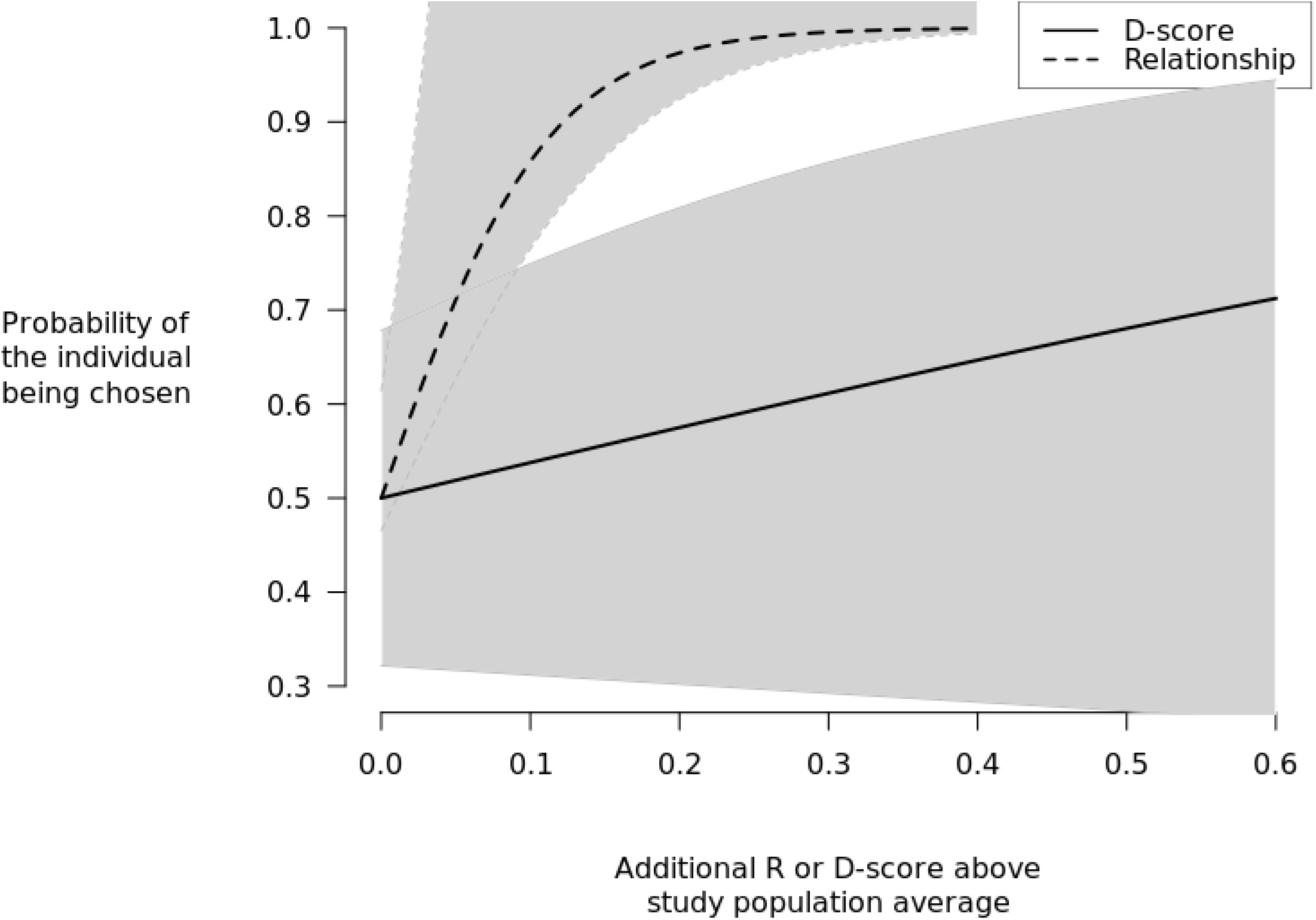
Plotted exploded logit model fit to MHC allele sharing data

We also tested whether MSAT *D*-score for pairs of participants in the FSR survey altered the probability of an individual being chosen, using the same logit model. A partial model, using only age difference and relationship as predictors, performed the same as the full model based on AIC score (**Table S3**), suggesting that MSAT *D*-score does not increase the probability of an individual being identified as visually similar. Using the full model, the MSAT plot reveals the same trend as the MHC allele sharing plot: for each 0.25 above average R score the coefficient of relationship is an important predictor of an individual being chosen (**Figure 2**). For each 0.1 *D*-score point above average, the probability of an individual being as visually similar increases by a few percentage points, though the estimate for the MSAT *D*-score effect is less accurate, as indicated by the wider confidence band. This result mirrors the positive correlation between MHC-DRB and MSAT *D*-score (**Figure S1**). However, the equal probability of the full and partial models combined with the large confidence band leads us to conclude that MSAT *D*-scores closer to one do not increase the probability of an individual being chosen as predictably as HLA-DRB *D*-score.

**Figure 2:**
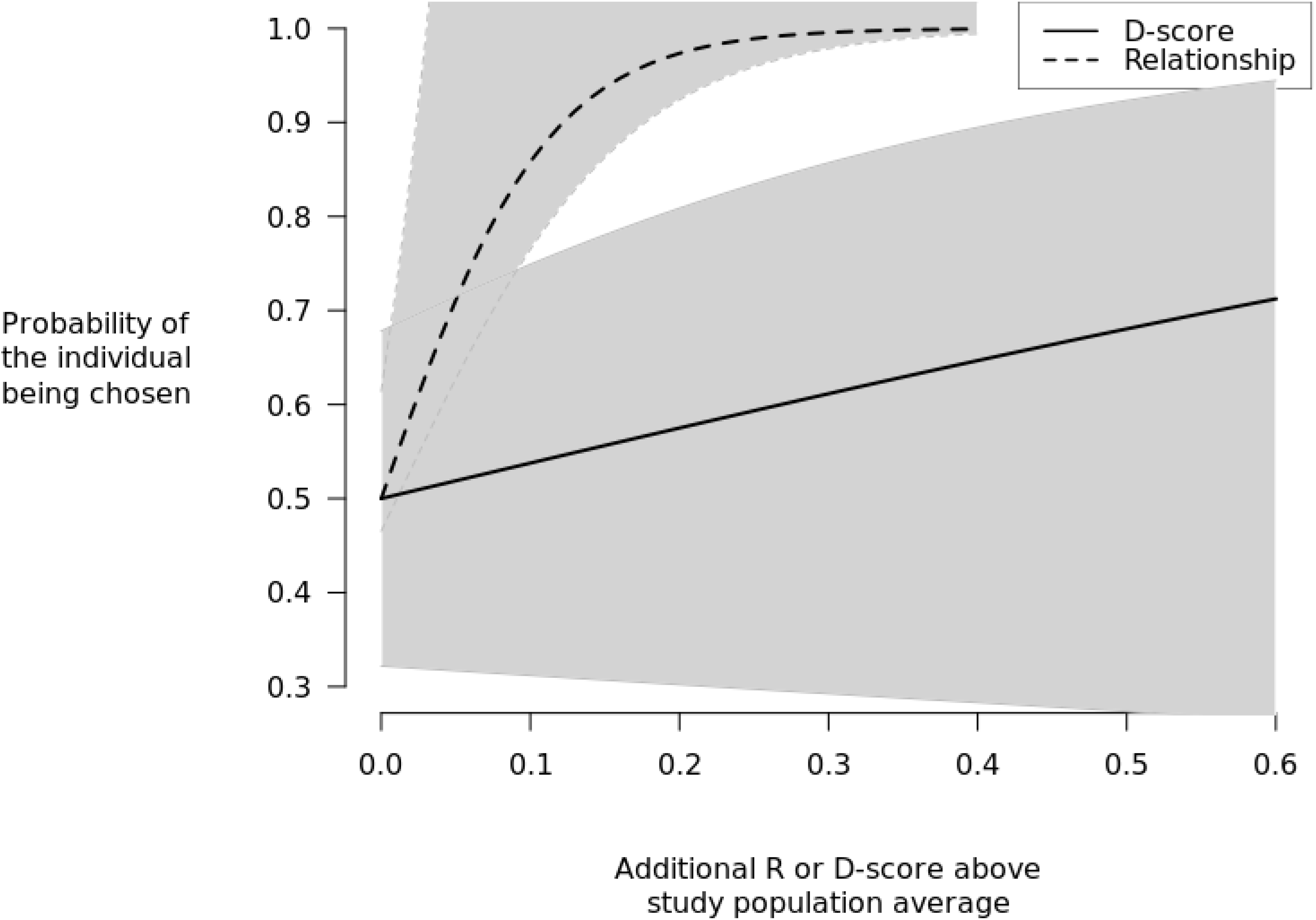
Plotted exploded logit model fit to MSAT allele sharing data

## 4 Discussion

Our results revealed that family relationship is a strong predictor of self-referent kin recognition. As expected, each participant’s first two FSR choices were always individuals within their own triad. Third choices varied among the members of each triad. Although the relationship curve plateaus because each participant has only two related individuals to choose from within their 68 options, the *D*-scores are generally greater between the participants and their three choices when compared to those not chosen. More specifically, as the *D*-score between two individuals increases, the probability of an individual being chosen continues to increase, even outside each relatedness triad. This means that greater HLA-DRB allele sharing increases the probability that an individual will be selected as “more similar” to the participant, even when the individual is not directly related.

The overall HLA-DRB diversity in our study population was low and many unrelated individuals shared HLA-DRB alleles and haplotypes. For our study, however, this was ideal since we were able to observe how an increase in HLA-DRB *D*-score above population average increased the probability of an individual being chosen, suggesting that individuals are able to distinguish their own kin and non-kin with greater HLA similarity and can differentiate between subtle differences in HLA genotype. A study with no HLA sharing between unrelated individuals could not have revealed the same insights.

Although increases in MSAT *D*-score also reflected an increase in the probability of an individual being chosen, the confidence intervals did not follow a strong positive trend, indicating that overall genetic similarity does not help distinguish kin and non-kin as strongly as HLA-DRB similarity. As noted above, HLA-DRB scores significantly correlated with MSAT *D*-scores in our dataset (p= 1.45E-51)(**Figure S1**), supporting previous conclusions that MHC similarity might be a reflection of overall genetic similarity [35].

Genetic studies of chromosomal anomalies have found predictable relationships between facial traits, such as nasal bridge and eyebrow shape or dental features, and microdeletions or translocations [9, 24] Facial similarity and perception of self-resemblance in the present study is likely unrelated to genetic and chromosomal abnormalities. Therefore, the feasibility of finding a positive relationship between genes and facial features is supported by previous studies of phenotype and chromosomal anomalies. While some research suggests that craniofacial morphology is at least partly due to genetic factors [5], the direct relationship between facial morphology and genes remains controversial [10].

HLA allelic similarity may manifest in facial cues that increase the probability of identification of self-resemblance in humans. Our results indicate that other genetic markers, such as MSATs and mtDNA, do not have the same impact. Future work should include HLA class I alleles and other nuclear genetic markers to clarify if specific HLA loci or haplotypes and/or other genes are most important in visual kin recognition. As expected, mtDNA identity was observed among biologically related females. Research on mtDNA haplogroups have shown a relationship between mtDNA and facial features. For example, mtDNA haplogroups of East African origin occur more frequently in females with relatively longer faces [21]. Unfortunately, for our study, mitochondrial haplotypes were not sufficiently different between unrelated participants to test our hypothesis using mtDNA. If we had found greater mitochondrial variation among our participants, we might have been able to determine if perceived self-resemblance is correlated with sharing of mtDNA haplotypes.

The use of protocols that link DNA data to an individual’s identity (i.e., DNA profiling) have been the foundation of forensic investigations for decades. Currently, efforts to develop panels of genetic markers for DNA-phenotyping are increasing [40]. Forensic scientists can now survey the genomic backgrounds of individuals, in the context of genetic variation shared by specific populations, to predict hair and/or eye color phenotypes and also ancestry [28]. In these studies, whole genomes are examined and identification of locus-specific factors is not the object. Our results suggest that some genetic markers, such as HLA-DRB, may have significant influence on phenotype. If this is the case, a large scale survey of HLA and facial feature morphology may not only yield valuable insight for evolutionary biologists, but also forensic scientists.

Based on our extensive literature review, previous studies in humans examining the relationship between self-resemblance, cooperative behavior and/or trust have not incorporated any real measure of genetic similarity between related or unrelated individuals. Similarly, research testing unrelated judge’s perceptions of facial similarity among relatives has also not accounted for actual genetic similarity in the relatives. As we have seen here, some mother/daughter, and even sister, pairs are 100% genetically identical rather than just 50%, as predicted by Mendelian inheritance.

Perhaps some inconsistencies in results from previous studies described above reflect the fact that there may be more, or less, genetic similarity between related individuals involved in experiments. Children sharing more HLA alleles with their mother, for example, may be perceived as less similar to their fathers if the father/child pair shares only 50% of their alleles. Contrastingly, children sharing more HLA alleles with their father might be viewed as less similar to their mothers. It would be interesting to repeat earlier experiments evaluating perceived similarities in close relatives and incorporate HLA-DRB genotyping.

In addition to olfaction, many nonhuman species also rely upon visual signals for kin recognition [38]. Studies of zebrafish, for example, have shown that family-specific morphometry and pigmentation patterns are all influenced by MHC genotype [26]. These visual cues are useful signals for kin recognition in zebrafish [26].

Given our reliance on vision, it would be advantageous for genes to help communicate similarity via facial cues. Indeed, the ability to confirm relatedness and allele sharing through self-resemblance would facilitate preferential behavior toward kin and, in turn, potentially increase the frequency of those shared alleles. In the future, it would also be interesting to control for kin familiarity to determine how much self-resemblance is distinct from familiarity in humans. In our species, there are many unanswered questions regarding the familiarity, self-resemblance, kin selection and mate-choice. The results of our study suggest that direct measures of genetic similarity may provide valuable insight into these long-standing questions.

## Acknowledgements

We thank Shane J. Macfarlan and Elizabeth Cashdan for helpful comments on this manuscript and Jeremy Koster for pointing us towards the exploded logit model.

