## Supplemental Tables and Figures for "Genetic similarity and facial cues for kin recognition in humans"

### Supplementary Material

#### Figures

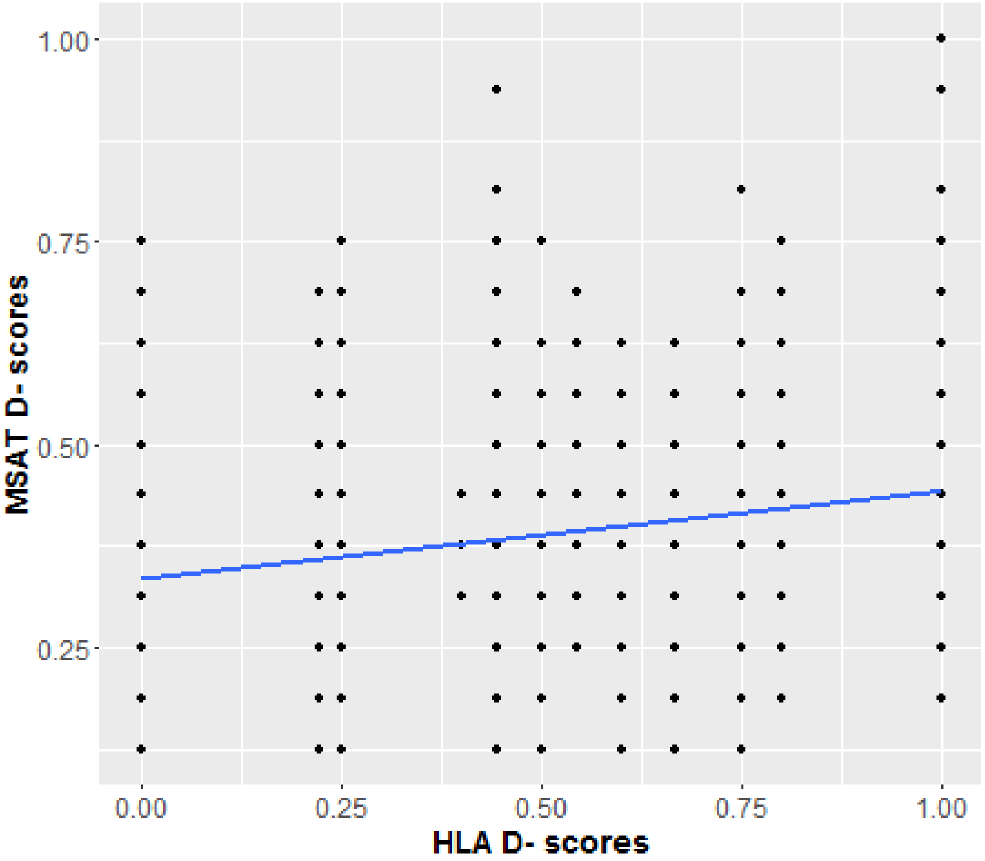


Figure S1: HLA $D$-score and MSAT $D$-score regression. R-squared $= 0.047$, p-value $= 0.45E-51$

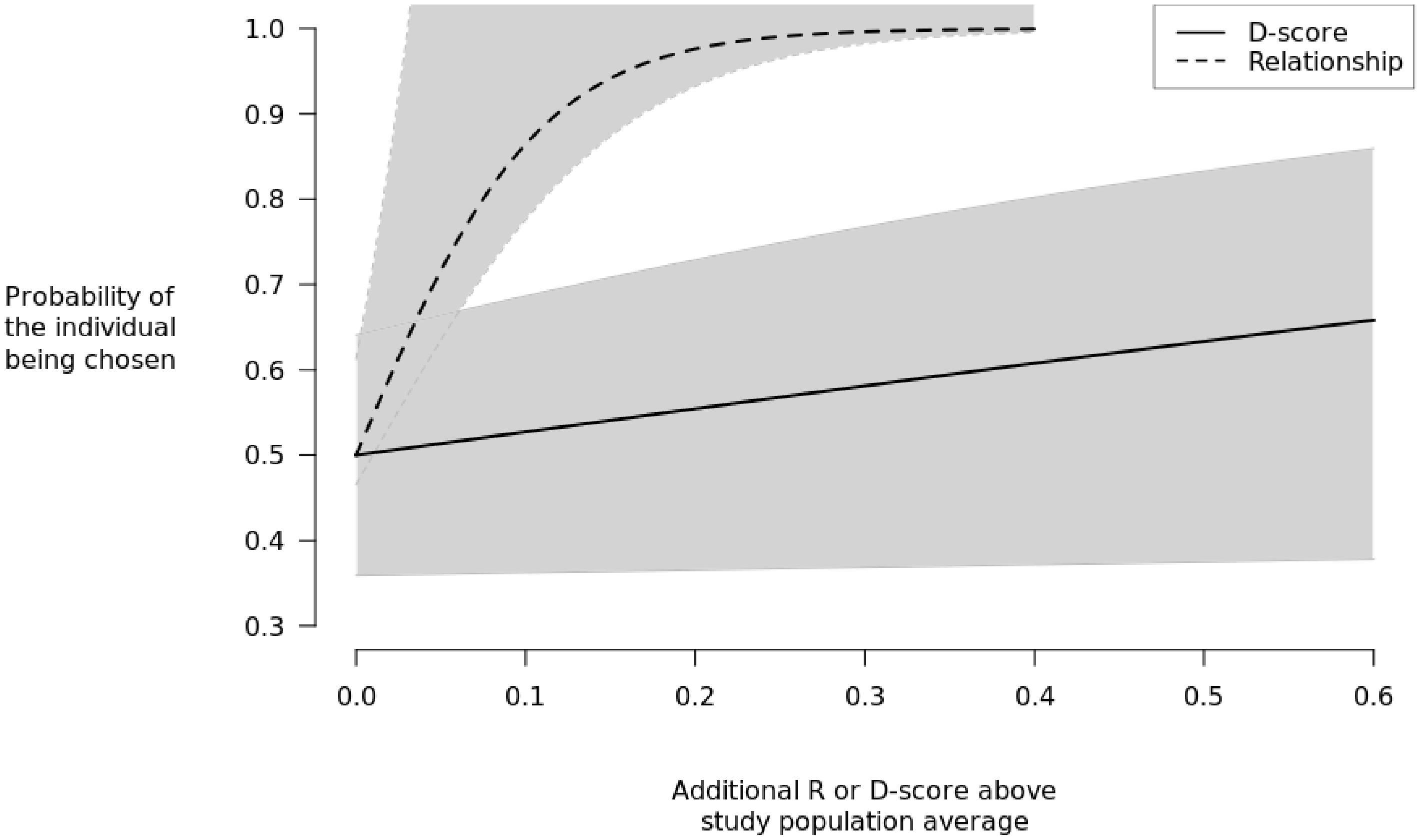


Figure S2: Plotted exploded logit model fit to serotype allele sharing data

### Tables

Table S1: Three variations of the exploded logit model fit to MHC data, model estimates, standard errors, and AIC scores. Full (age, D-score, and relationship): $u_{ij} = A_{ij}\beta_A + D_{ij}\beta_D + R_{ij}\beta_R$ Partial 1 (age and D-score): $u_{ij} = A_{ij}\beta_A + D_{ij}\beta_D$ Partial 2 (age and relationship): $u_{ij} = A_{ij}\beta_A + R_{ij}\beta_R$

| | **Full** | **Full standard error** | **Partial 1** | **Partial 1 standard error** | **Partial 2** | **Partial 2 standard error** |
|---|---|---|---|---|---|---|
| **Age difference** | -0.0015 | 0.0062 | -0.0065 | 0.0043 | -0.0018 | 0.0062 |
| **D-score** | 1.2346 | 0.4232 | 3.505 | 0.3025 | – | – |
| **R** | 18.2507 | 2.3824 | – | – | 18.69 | 2.366 |
| **AIC** | 687 | | 1599 | | 694 | |

Table S2: Three variations of the exploded logit model fit to serotype data, model estimates, standard errors, and AIC scores. Full (age, D-score, and relationship): $u_{ij} = A_{ij}\beta_A + D_{ij}\beta_D + R_{ij}\beta_R$ Partial 1 (age and D-score): $u_{ij} = A_{ij}\beta_A + D_{ij}\beta_D$ Partial 2 (age and relationship): $u_{ij} = A_{ij}\beta_A + R_{ij}\beta_R$

| | **Full** | **Full standard error** | **Partial 1** | **Partial 1 standard error** | **Partial 2** | **Partial 2 standard error** |
|---|---|---|---|---|---|---|
| **Age difference** | -0.0014 | 0.0062 | -0.0061 | 0.0044 | -0.0018 | 0.0062 |
| **D-score** | 1.0937 | 0.4789 | 2.3618 | 0.3403 | – | – |
| **R** | 18.5431 | 2.3752 | – | – | 18.69 | 2.366 |
| **AIC** | 691 | | 1692 | | 694 | |

Table S3: Three variations of the exploded logit model fit to serotype data, model estimates, standard errors, and AIC scores. Full (age, D-score, and relationship): $u_{ij} = A_{ij}\beta_A + D_{ij}\beta_D + R_{ij}\beta_R$ Partial 1 (age and D-score): $u_{ij} = A_{ij}\beta_A + D_{ij}\beta_D$ Partial 2 (age and relationship): $u_{ij} = A_{ij}\beta_A + R_{ij}\beta_R$

| | **Full** | **Full standard error** | **Partial 1** | **Partial 1 standard error** | **Partial 2** | **Partial 2 standard error** |
|---|---|---|---|---|---|---|
| **Age difference** | -0.0012 | 0.0062 | 8.20E-04 | 0.0048 | -0.0018 | 0.0062 |
| **D-score** | 1.5114 | 0.9857 | 1.3E-01 | 0.6558 | – | – |
| **R** | 18.003 | 2.4081 | – | – | 18.69 | 2.366 |
| **AIC** | 694 | | 1275 | | 694 | |

Table S4: The **FSR data table** reports the first, second and third choices of each member of the Sample group in the FSR survey and the HLA, MSAT and Serotype $D$-scores between the participant and each choice, including the average score between the participant and all of the other individuals they did not choose.

| **Part.** | **Part. Age** | **C1** | **MHC** $D$**-score** | **Sero.** $D$**-score** | **MSAT** $D$**-score** | **C2** | **MHC** $D$**-score** | **Sero.** $D$**-score** | **MSAT** $D$**-score** | **C3** | **MHC** $D$**-score** | **Sero.** $D$**-score** | **MSAT** $D$**-score** |
|---|---|---|---|---|---|---|---|---|---|---|---|---|---|
| 1A | 19 | 1C | 1.0000 | 1.0000 | 0.6250 | 1B | 1.0000 | 1.0000 | 0.8750 | 13B | 0.2500 | 0.6667 | 0.3125 |
| 1B | 17 | 1A | 1.0000 | 1.0000 | 0.8750 | 1C | 1.0000 | 1.0000 | 0.7500 | 11A | 0.6667 | 1.0000 | 0.5000 |
| 1C | 15 | 1A | 1.0000 | 1.0000 | 0.6250 | 1B | 1.0000 | 1.0000 | 0.7500 | 3B | 0.4000 | 0.6667 | 0.3125 |
| 2A | 18 | 2B | 0.7500 | 1.0000 | 0.7500 | 2C | 0.5000 | 0.5714 | 0.6250 | 12B | 0.2222 | 0.5714 | 0.1875 |
| 2B | 46 | 2A | 0.7500 | 1.0000 | 0.7500 | 2C | 0.5000 | 0.5714 | 0.5625 | 21B | 0.2500 | 0.5714 | 0.5000 |
| 2C | 20 | 2B | 0.5000 | 0.5714 | 0.5625 | 2A | 0.5000 | 0.5714 | 0.6250 | 5B | 0.6667 | 1.0000 | 0.3750 |
| 3A | 22 | 3C | 0.4444 | 0.5714 | 0.7500 | 3B | 0.0000 | 0.0000 | 0.6875 | 5B | 0.4444 | 0.5714 | 0.3125 |
| 3B | 50 | 3C | 0.5454 | 0.3333 | 0.6875 | 3A | 0.0000 | 0.0000 | 0.6875 | 5C | 0.0000 | 0.0000 | 0.4375 |
| 3C | 19 | 3B | 0.5454 | 0.3333 | 0.6875 | 3A | 0.4444 | 0.5714 | 0.7500 | 10C | 0.8000 | 0.7500 | 0.2500 |
| 4A | 83 | 4B | 0.5000 | 0.6667 | 0.6875 | 4C | 0.5000 | 0.6667 | 0.6875 | 8B | 0.2222 | 0.5714 | 0.3125 |
| 4B | 51 | 4A | 0.5000 | 0.6667 | 0.6875 | 4C | 0.7500 | 1.0000 | 0.5625 | 14C | 0.5000 | 0.6667 | 0.4375 |
| 4C | 58 | 4B | 0.7500 | 0.6667 | 0.5625 | 4A | 0.5000 | 1.0000 | 0.6875 | 11C | 0.5000 | 0.6667 | 0.3750 |
| 5A | 20 | 5C | 1.0000 | 1.0000 | 0.6250 | 5B | 0.4444 | 0.6667 | 0.8125 | 18A | 0.0000 | 0.3333 | 0.4375 |
| 5B | 18 | 5C | 0.4444 | 0.6667 | 0.6875 | 5A | 0.4444 | 0.6667 | 0.8125 | 20A | 0.8000 | 1.0000 | 0.3750 |
| 5C | 47 | 5A | 1.0000 | 1.0000 | 0.6250 | 5B | 0.4444 | 0.6667 | 0.6875 | 18A | 0.0000 | 0.3333 | 0.3750 |
| 6A | 80 | 6C | 0.5000 | 0.2857 | 0.5625 | 6B | 0.4444 | 0.3333 | 0.5000 | 20C | 0.2500 | 0.4000 | 0.1875 |
| 6B | 67 | 6C | 0.2222 | 0.5714 | 0.4375 | 6A | 0.4444 | 0.3333 | 0.5000 | 18C | 0.4444 | 0.6667 | 0.3125 |
| 6C | 61 | 6B | 0.2222 | 0.5714 | 0.4375 | 6A | 0.5000 | 0.2857 | 0.5625 | 22B | 0.0000 | 0.2500 | 0.3750 |
| 7A | 24 | 7C | 0.7500 | 0.6667 | 0.5000 | 7B | 0.7500 | 0.6667 | 0.6250 | 6B | 0.4444 | 0.6667 | 0.4375 |
| 7B | 22 | 7C | 1.0000 | 1.0000 | 0.6875 | 7A | 0.7500 | 0.6667 | 0.6250 | 4B | 0.5000 | 1.0000 | 0.5000 |
| 7C | 53 | 7B | 1.0000 | 1.0000 | 0.6875 | 7A | 0.7500 | 0.6667 | 0.5000 | 9A | 0.5000 | 0.5714 | 0.5000 |
| 8A | 23 | 8C | 0.8000 | 0.5000 | 0.5625 | 8B | 0.8000 | 0.7500 | 0.6250 | 13A | 0.4444 | 0.6667 | 0.6250 |
| 8B | 56 | 8A | 0.8000 | 0.7500 | 0.6250 | 8C | 1.0000 | 0.7500 | 0.8125 | 11C | 0.2222 | 0.2857 | 0.5000 |
| 8C | 79 | 8A | 0.8000 | 0.5000 | 0.5625 | 8B | 1.0000 | 0.7500 | 0.8125 | 7C | 0.4444 | 0.5714 | 0.2500 |
| 9A | 24 | 9B | 0.4444 | 0.5000 | 0.5625 | 9C | 0.4444 | 0.5000 | 0.6250 | 18C | 0.5000 | 0.7500 | 0.4375 |
| 9B | 21 | 9C | 1.0000 | 1.0000 | 0.6250 | 9A | 0.4444 | 0.5000 | 0.5625 | 19C | 0.4444 | 0.5714 | 0.1875 |
| 9C | 55 | 9A | 0.4444 | 0.5000 | 0.6250 | 9B | 1.0000 | 1.0000 | 0.6250 | 12C | 0.6000 | 0.7500 | 0.5000 |
| 10A | 20 | 10C | 1.0000 | 1.0000 | 0.4375 | 10B | 1.0000 | 1.0000 | 0.7500 | 23C | 0.8000 | 1.0000 | 0.5000 |
| 10B | 50 | 10A | 1.0000 | 1.0000 | 0.7500 | 10C | 1.0000 | 1.0000 | 0.5625 | 22B | 0.8000 | 1.0000 | 0.5000 |
| 10C | 23 | 10A | 1.0000 | 1.0000 | 0.4375 | 10B | 1.0000 | 1.0000 | 0.5625 | 23A | 0.2222 | 0.5000 | 0.2500 |
| 11A | 20 | 11B | 0.4444 | 0.5714 | 0.5625 | 11C | 0.4444 | 0.5714 | 0.3750 | 19C | 0.4444 | 0.5714 | 0.5625 |
| 11B | 54 | 11C | 1.0000 | 0.5714 | 0.7500 | 11A | 0.4444 | 1.0000 | 0.5625 | 20A | 0.2222 | 0.5714 | 0.3750 |
| 11C | 22 | 11B | 1.0000 | 0.5714 | 0.7500 | 11A | 0.4444 | 1.0000 | 0.3750 | 23A | 0.5000 | 0.5714 | 0.1875 |
| 12A | 48 | 12B | 0.8000 | 1.0000 | 0.5625 | 12C | 0.8000 | 1.0000 | 0.6250 | 21B | 0.2222 | 0.5000 | 0.5000 |
| 12B | 19 | 12C | 1.0000 | 1.0000 | 0.5625 | 12A | 0.8000 | 1.0000 | 0.5625 | 13B | 0.2222 | 0.6667 | 0.1875 |
| 12C | 18 | 12B | 1.0000 | 1.0000 | 0.5625 | 12A | 0.8000 | 1.0000 | 0.6250 | 13A | 0.2222 | 0.6667 | 0.3750 |
| 13A | 18 | 13B | 0.7500 | 0.8000 | 0.3125 | 13C | 0.5000 | 1.0000 | 0.5625 | 15B | 0.2222 | 0.3333 | 0.2500 |
| 13B | 21 | 13C | 0.5000 | 0.8000 | 0.5625 | 13A | 0.7500 | 1.0000 | 0.3125 | 16C | 0.2500 | 0.5000 | 0.2500 |
| 13C | 23 | 13B | 0.5000 | 0.8000 | 0.5625 | 13A | 0.5000 | 0.8000 | 0.5625 | 19C | 0.5000 | 0.6667 | 0.3125 |
| 14A | 24 | 14B | 1.0000 | 1.0000 | 0.5625 | 14C | 0.5000 | 0.4000 | 0.5625 | 10A | 0.2222 | 0.3333 | 0.5625 |
| 14B | 55 | 14C | 0.5000 | 0.4000 | 0.6875 | 14A | 1.0000 | 1.0000 | 0.5625 | 8A | 0.4444 | 0.3333 | 0.3750 |
| 14C | 22 | 14B | 0.5000 | 0.4000 | 0.6875 | 14A | 0.5000 | 0.4000 | 0.5625 | 18B | 0.4444 | 0.5714 | 0.1875 |
| 15A | 46 | 15B | 0.4444 | 0.5714 | 0.6875 | 15C | 0.4444 | 0.5714 | 0.5625 | 9B | 0.4444 | 0.5714 | 0.5625 |
| 15B | 19 | 15C | 0.8000 | 0.7500 | 0.5625 | 15A | 0.4444 | 0.5714 | 0.6875 | 9A | 0.4444 | 0.5000 | 0.3125 |
| 15C | 18 | 15B | 0.8000 | 0.7500 | 0.5625 | 15A | 0.4444 | 0.5714 | 0.5625 | 18B | 0.8000 | 0.7500 | 0.3125 |
| 16A | 22 | 16C | 0.4444 | 1.0000 | 0.9375 | 16B | 0.4444 | 0.5000 | 0.6250 | 6B | 1.0000 | 0.8571 | 0.4275 |
| 16B | 55 | 16C | 0.5000 | 0.5000 | 0.5625 | 16A | 0.4444 | 0.5000 | 0.6250 | 15A | 0.5000 | 0.5714 | 0.3125 |
| 16C | 25 | 16B | 0.5000 | 0.6667 | 0.5625 | 16A | 0.4444 | 0.6667 | 0.9375 | 19A | 0.5000 | 0.8000 | 0.4375 |
| 17A | 52 | 17C | 0.4444 | 0.6667 | 0.6250 | 17B | 0.4444 | 0.6667 | 0.6250 | 4A | 0.2500 | 0.8000 | 0.3750 |
| 17B | 18 | 17C | 1.0000 | 1.0000 | 0.8125 | 17A | 0.4444 | 0.6667 | 0.6250 | 15B | 1.0000 | 1.0000 | 0.3750 |
| 17C | 19 | 17B | 1.0000 | 1.0000 | 0.8125 | 17A | 0.4444 | 0.6667 | 0.6250 | 9B | 1.0000 | 1.0000 | 0.4375 |
| 18A | 54 | 18B | 1.0000 | 1.0000 | 0.6875 | 18C | 0.4444 | 0.5714 | 0.6250 | 20A | 0.6000 | 0.7500 | 0.5000 |
| 18B | 22 | 18A | 1.0000 | 1.0000 | 0.6875 | 18C | 0.4444 | 0.5714 | 0.7500 | 15C | 0.8000 | 0.7500 | 0.3125 |
| 18C | 19 | 18A | 0.4444 | 0.5714 | 0.6250 | 18B | 0.4444 | 0.5714 | 0.7500 | 6B | 0.4444 | 0.6667 | 0.3125 |
| 19A | 25 | 19B | 0.8571 | 0.4000 | 0.5000 | 19C | 1.0000 | 1.0000 | 0.3750 | 15A | 0.7500 | 0.6667 | 0.3750 |
| 19B | 56 | 19C | 0.8571 | 0.8000 | 0.5625 | 19A | 0.8571 | 0.8000 | 0.5000 | 11B | 0.5000 | 0.8000 | 0.3125 |
| 19C | 53 | 19B | 0.8571 | 0.8000 | 0.5625 | 19A | 1.0000 | 1.0000 | 0.3750 | 11C | 0.5000 | 0.6667 | 0.3750 |
| 20A | 71 | 20B | 1.0000 | 1.0000 | 0.5000 | 20C | 0.4444 | 0.6667 | 0.4375 | 17A | 0.0000 | 0.3333 | 0.2500 |
| 20B | 49 | 20C | 0.4444 | 0.6667 | 0.6250 | 20A | 1.0000 | 1.0000 | 0.5000 | 14A | 0.2222 | 0.3333 | 0.3125 |
| 20C | 18 | 20B | 0.4444 | 0.6667 | 0.6250 | 20A | 0.4444 | 0.6667 | 0.4375 | 17B | 0.0000 | 0.3333 | 0.3125 |
| 21A | 42 | 21B | 0.5000 | 0.5714 | 0.6250 | 21C | 0.2500 | 0.5714 | 0.7500 | 10A | 0.4444 | 0.5714 | 0.5625 |
| 21B | 40 | 21C | 0.7500 | 1.0000 | 0.8125 | 21A | 0.5000 | 0.5714 | 0.6250 | 22A | 0.2222 | 0.5000 | 0.4375 |
| 21C | 67 | 21B | 0.7500 | 1.0000 | 0.8125 | 21A | 0.2500 | 0.5714 | 0.7500 | 10C | 0.2222 | 0.5000 | 0.5625 |
| 22A | 45 | 22B | 1.0000 | 1.0000 | 0.5000 | 22C | 1.0000 | 1.0000 | 0.6250 | 2A | 0.2222 | 0.5714 | 0.1875 |
| 22B | 66 | 22A | 1.0000 | 1.0000 | 0.5000 | 22C | 1.0000 | 1.0000 | 0.6250 | 3C | 0.8000 | 0.7500 | 0.2500 |
| 22C | 69 | 22A | 1.0000 | 1.0000 | 0.6250 | 22B | 1.0000 | 1.0000 | 0.6250 | 21C | 0.2222 | 0.5000 | 0.3750 |
| 23A | 57 | 23B | 0.4444 | 0.5000 | 0.5000 | 23C | 0.4444 | 0.5000 | 0.2500 | 21B | 0.7500 | 1.0000 | 0.4375 |
| 23B | 23 | 23A | 0.4444 | 0.5000 | 0.5000 | 23C | 1.0000 | 1.0000 | 0.4375 | 10A | 0.8000 | 1.0000 | 0.4375 |
| 23C | 25 | 23B | 1.0000 | 1.0000 | 0.4375 | 23A | 0.4444 | 0.5000 | 0.2500 | 14B | 0.4444 | 0.3333 | 0.4375 |

### Methods

```
# DATA PREP
#setwd(wd)

# dscore is an n x n matrix of D−scores
# between individual i and j
dscore <− read.csv( "file", header=TRUE, row.names=1 )

# create the delta (choice) matrix for each individual
# delta is a list per individual that contains a 4 x n matrix
# of ones and zeros; each row represents the choices available
# before each decision, where ones indicate choices available,
# and zeros not available corresponding to an individual chosen
# in the previous decision; the numerical position in the ID
# number of the individuals
d <− read.csv( "matrix_choices.csv", header=TRUE )
n <− dim(d)[1]
# delta <− lapply(1:n, function(z){
        # ans <− matrix( rep( 1, n∗(n−1)), nrow=(n−1) )
        # ch1 <− d[z,"Choice_1_ID"]
        # ch2 <− d[z,"Choice_2_ID"]
        # ch3 <− d[z,"Choice_3_ID"]
        # ans[2, ch1] <− 0
        # ans[3, c(ch1, ch2) ] <− 0
        # ans[4, c(ch1, ch2, ch3)] <− 0
        # for( i in 5:(n−1) ) ans[i, c(ch1, ch2, ch3)] <− 0
        # ans
        # })
delta <− lapply(1:n, function(z){
        ans <− matrix( rep( 1, n∗4), nrow=4 )
        ch1 <− d[z,"Choice_1_ID"]
        ch2 <− d[z,"Choice_2_ID"]
        ch3 <− d[z,"Choice_3_ID"]
        ans[2, ch1] <− 0
        ans[3, c(ch1, ch2) ] <− 0
        ans[4, c(ch1, ch2, ch3)] <− 0
        ans
        })
names(delta) <− paste0("n",1:n)
```

```
# R is an n x n matrix of Relatedness measures
# between individual i and j
R <- read.csv( "matrix_R.csv", header=TRUE, row.names=1 )

# age difference matrix
# agediff is a n x n matrix of age differences between
# individual i and j
agediff <- matrix( rep(NA,n*n), nrow=n)
for( i in 1:n ){
	for( j in 1:n ){
		agediff[i,j] <- d[i,"ParticipantAge"] - d[j,"ParticipantAge"]
	}
}



# check relationship between explanatory
# variables
cor( dscore, R)

# -----------
# FULL MODEL DSCORE AND R
nllik_mDscore_R <- function ( x ){

	# x is a 1x3 vector of model parameters
	# which includes coefficient on age differences,
	# coefficient on D score, coefficient on relatedness
	# x[1] is coefficient on age
	# x[2] is coefficient on D score
	# x[3] is coefficient on relatedness
	#x <- c(0.1, 0.2, 0.3)

	# the coefficients and explanatory variables
	# specify the utility or preference for a particular
	# choice

	# agediff is a n x n matrix of age differences between
	# individual i and j

	# dscore is an n x n matrix of D-scores
```

```
    # between individual i and j

    # R is an n x n matrix of Relatedness measures
    # between individual i and j

    # for any individual i, the preferences for choice j
    # is going to be a linear combination:
    # lets define a preference function, u, which
    # is a n x n matrix that defines the preference of
    # individual i for choice j

    u <- matrix( rep(NA, n*n), nrow=n )
    for( i in 1:n ){
        for ( j in 1:n ){
        u[i,j] <- abs(agediff[i,j]) * x[1] + dscore[i,j] * x[2] + R[i,j] * x[3]
            }
        }

    # delta is computed for each individual i
    # where each row corresponds to the choice
    # of each item, 0 for if the item is already chosen
    # previously, and 1 if not (or tied)

    # therefore delta is a list of length n
    # containing a (n-1) x n matrix of 1s and 0s
    # corresponding to individual i's (n-1) choices

    # for each individual i lets calculate the likelihood of their choices
    # given the model
    logL <- rep(NA,n)
    for( i in 1:n ){
        ch <- as.numeric(d[i,c("Choice_1_ID","Choice_2_ID","Choice_3_ID") ])
        oth <- seq(1,n,1)[-i] # this leaves out individual i
        logL[i] <- sum( sapply( 1:3, function(j) u[i,ch[j]] - log( sum( delta[[i]][j,oth]*exp( u[i,oth] ) ) ) ) )
        #logL[i] <- sum( u[i,oth] ) - sum( sapply( 1:4, function(j){ log( sum( delta[[i]][j,oth]*exp( u[i,oth] ) ) ) }
          ) )
                }

    -sum(logL)
}
```

```
x.inits <- c(0,1,18)
m1 <- optim( x.inits, nllik_mDscore_R, method="Nelder-Mead", hessian=TRUE, control=list(trace=TRUE, maxit
    =2000))

m1$convergence # value of zero mean successful convergence
m1$par # look at estimated parameters
m1$val # negative log-likelihood
m1parse <- diag( sqrt( solve( m1$hessian ) ) ) # estimate standard error

# ---------------
# PARTIAL MODEL DSCORE
nllik_mDscore <- function ( x ){

        u <- matrix( rep(NA, n*n), nrow=n )
        for( i in 1:n ){
                for ( j in 1:n ){
                u[i,j] <- abs(agediff[i,j]) * x[1] + dscore[i,j] * x[2]
                        }
                }

        logL <- rep(NA,n)
        for( i in 1:n ){
                ch <- as.numeric(d[i,c("Choice_1_ID","Choice_2_ID","Choice_3_ID") ])
                oth <- seq(1,n,1)[-i] # this leaves out individual i
                logL[i] <- sum( sapply( 1:3, function(j) u[i,ch[j]] - log( sum( delta[[i]][j,oth]*exp( u[i,oth] ) ) ) ) )
                                }
        -sum(logL)
}

x.inits <- c(0,1)
m2 <- optim( x.inits, nllik_mDscore, method="Nelder-Mead", hessian=TRUE, control=list(trace=TRUE, maxit=2000)
    )

m2$convergence # value of zero mean successful convergence
m2$par # look at estimated parameters
m2$val # negative log-likelihood
m2parse <- diag( sqrt( solve( m2$hessian ) ) ) # estimate standard error
```

```
# -----------
# PARTIAL MODEL JUST R
nllik_R <- function ( x ){

    u <- matrix( rep(NA, n*n), nrow=n )
    for( i in 1:n ){
        for ( j in 1:n ){
        u[i,j] <- abs(agediff[i,j]) * x[1] + R[i,j] * x[2]
            }
        }

    logL <- rep(NA,n)
    for( i in 1:n ){
        ch <- as.numeric(d[i,c("Choice_1_ID","Choice_2_ID","Choice_3_ID") ])
        oth <- seq(1,n,1)[-i] # this leaves out individual i
        logL[i] <- sum( sapply( 1:3, function(j) u[i,ch[j]] - log( sum( delta[[i]][j,oth]*exp( u[i,oth] ) ) ) ) )
                }
    -sum(logL)
}

x.inits <- c(0,18.693)
m3 <- optim( x.inits, nllik_R, method="Nelder-Mead", hessian=TRUE, control=list(trace=TRUE, maxit=2000))

m3$convergence # value of zero mean successful convergence
m3$par # look at estimated parameters
m3$val # negative log-likelihood
m3parse <- diag( sqrt( solve( m3$hessian ) ) ) # estimate standard error

# Results table
aicm <- function( m ) format( 2*length(m$par) - 2*(-m$value), digits=2)

m1pars <- c(format( m1$par, digits=2),aicm(m1))
m1se <- c(format(m1parse,digits=2),"--")
m2pars <- c(format( m2$par, digits=2),"--",aicm(m2))
m2se <- c(format(m2parse,digits=2),"--","--")
m3pars <- c(format( m3$par[1], digits=2),"--",format( m3$par[2], digits=4),aicm(m3))
m3se <- c(format(m3parse[1],digits=2),"--",format(m3parse[2],digits=4),"--")

res <- data.frame( m1pars, m1se, m2pars, m2se, m3pars, m3se, row.names=c("age.diff", "dscore", "R","AIC") )
```

```
res

#save.image( file="dscoreAnalysis.rdata" )

# Plot effects

utilmean <- function( dB, RB ) {
        exp( dB * mean(unlist(dscore)) + RB  * mean( unlist(R) ) )
        }

utilnew <- function( dB, RB, add_ds=0, add_R=0 ) {
        exp( dB * (mean(unlist(dscore)) + add_ds ) + RB * (mean( unlist(R) ) + add_R ) )
        }

add_ds <- seq(0,0.6,0.01)
probChDscore <- utilnew( dB=m1$par[2], RB=m1$par[3], add_ds=add_ds ) / ( utilmean( dB=m1$par[2], RB=m1$par
    [3] ) + utilnew( dB=m1$par[2], RB=m1$par[3], add_ds=add_ds ) )
probChDscore_lb <- utilnew( dB=(m1$par[2]-2*m1parse[2]), RB=m1$par[3], add_ds=add_ds ) / ( utilmean( dB=
    m1$par[2], RB=m1$par[3] ) + utilnew( dB=(m1$par[2]-2*m1parse[2]), RB=m1$par[3], add_ds=add_ds ) )
probChDscore_ub <- utilnew( dB=(m1$par[2]+2*m1parse[2]), RB=m1$par[3], add_ds=add_ds ) / ( utilmean( dB=
    m1$par[2], RB=m1$par[3] ) + utilnew( dB=(m1$par[2]+2*m1parse[2]), RB=m1$par[3], add_ds=add_ds ) )

add_R <- seq(0,0.4,0.01)
probChR <- utilnew( dB=m1$par[2], RB=m1$par[3], add_R=add_R ) / ( utilmean( dB=m1$par[2], RB=m1$par[3] ) +
     utilnew( dB=m1$par[2], RB=m1$par[3], add_R=add_R ) )
probChR_lb <- utilnew( dB=m1$par[2], RB=(m1$par[3]-2*m1parse[3]), add_R=add_R ) / ( utilmean( dB=m1$par[2],
     RB=m1$par[3] ) + utilnew( dB=m1$par[2], RB=(m1$par[3]-2*m1parse[3]), add_R=add_R ) )
probChR_ub <- utilnew( dB=m1$par[2], RB=(m1$par[3]+2*m1parse[3]), add_R=add_R ) / ( utilmean( dB=m1$par
    [2], RB=m1$par[3] ) + utilnew( dB=m1$par[2], RB=(m1$par[3]-2*m1parse[3]), add_R=add_R ) )



par( mar=c(7,10,1,1), las=1)
plot( x=add_ds, probChDscore, type="l", axes=FALSE, lwd=2, ylim=c(0.3,1), ylab="", xlab="")
axis(1)
axis(2)
mtext( "Additional R or D-score above\nstudy population average", side=1, line=5)
mtext( "Probability of the\n individual\nbeing chosen", side=2, line=10, adj=0)
legend( "topright", lty=1:2, lwd=2, legend=c("D-score", "Relationship"), box.lwd=0)
polygon( x=c(add_R,rev(add_R)), y=c(probChR_ub,rev(probChR_lb)), col="lightgray", border="white")
```

```
polygon( x=c(add_ds,rev(add_ds)), y=c(probChDscore_ub,rev(probChDscore_lb)), col="lightgray", border="white")
lines( x=add_R, y=probChR_lb, lty=2, lwd=1, col="gray")
lines( x=add_R, y=probChR_ub, lty=2, lwd=1, col="gray")
lines( x=add_R, y=probChR, lty=2, lwd=2 )
lines( x=add_ds, y=probChDscore_lb, lty=1, lwd=1, col="gray")
lines( x=add_ds, y=probChDscore_ub, lty=1, lwd=1, col="gray")
lines( x=add_ds, y=probChDscore, lty=1, lwd=2 )
```
